# In Silico Genetics: Identification of pathogenic nsSNPs in human *STAT3* gene associated with Job’s syndrome

**DOI:** 10.1101/545657

**Authors:** Mujahed I. Mustafa, Abdelrahman H. Abdelmoneim, Nafisa M. Elfadol, Thwayba A. Mahmoud, Mohamed A. Hassan

**Affiliations:** Department of Biochemistry, University of Bahri, Sudan; Department of Biotechnology, Africa city of Technology, Sudan

**Keywords:** Autosomal dominant hyper-IgE syndrome *(AD-HIES)*, Job’s syndrome, *STAT3* gene, in silico analysis, *diagnostic markers*

## Abstract

**Background:** Autosomal dominant hyper-IgE syndrome (AD-HIES) or Job’s syndrome is a rare immunodeficiesncy disease that classically presents in early childhood, characterized by eczematoid dermatitis, characteristic facies, pneumatoceles, hyperextensibility of joints, multiple bone fractures, scoliosis, atopic dermatitis and elevated levels of serum IgE (>2000 IU/ml). The term Autosomal dominant hyper-IgE syndrome has primarily been associated with mutations in STAT3 gene, Located in human chromosome 17q21.

**Methods:** The human STAT3 gene was investigated in dbSNP/NCBI, 962 SNPs were Homo sapiens; of which 255 were missense SNPs. This selected for in silico analysis by multiple in silico tools to investigate the effect of SNPs on *STAT3* protein’s structure and function.

**Result:** Eleven novel mutations out of 255 nsSNPs that are found to be deleterious effect on the STAT3 structure and function.

**Conclusion:** A total of eleven novel nsSNPs were predicted to be responsible for the structural and functional modifications of STAT3 protein. The newly recognized genetic cause of the hyper-IgE syndrome affects complex, compartmentalized somatic and immune regulation. This study will opens new doors to facilitate the development of novel diagnostic markers for associated diseases.

## Introduction

Autosomal dominant hyper-IgE syndrome (AD-HIES) or Job’s syndrome is a rare immunodeficiesncy disease that classically presents in early childhood, characterized by eczematoid dermatitis, characteristic facies, pneumatoceles, hyperextensibility of joints, multiple bone fractures, scoliosis, atopic dermatitis and elevated levels of serum IgE (>2000 IU/ml).[1–11] Job’s syndrome was frst described by Davis et al. in 1966[12] Since then, patients from different countries are being increasingly recognized. [2, 6, 13–18]

The treatment for Hyper-IgE syndromes is mainly to control infection, skin care with Cyclosporine A, if necessary, should be done as early as possible hematopoietic stem cell transplantation.[3, 10, 19] The term Autosomal dominant hyper-IgE syndrome has primarily been associated with mutations in *STAT3* gene. [5, 8] Located in human chromosome 17q21, the other genes that have been implicated in HIES include *TYK2*[20, 21] and *DOCK8*[22–24] *STAT3* plays a vital role in signal transduction induced by many cytokines (IL-6, IL-10, IL-17, IL-21, and IL-22).[25, 26] Mutations lead to disruption of STAT3-dependent pathways, which are crucial for signaling of many cytokines, including IL-6 and IL-10.[27]

*STAT3* also plays important roles in multiple aspects of cancer aggressiveness including migration, invasion, survival, self-renewal, angiogenesis, and tumor cell immune evasion by regulating the expression of multiple downstream target genes.[28, 29] Some studies show that loss-of-function *STAT3* alleles were shown to be dominant-negative, led to a more detailed description of the immunologic phenotype of this primary immunodeficiency, with the demonstration of a deficiency of IL-17A- and IL-22-producing T cells.[5, 30, 31] While other study reveals a pattern of STAT3-associated gene expression specific to basal-like breast cancers in human tumors.[32]. [28, 29] therefore, the treatment for Hyper-IgE syndromes is crucial to prevent a serious complication in AD-HIES patients as cystic lung disease[33] However, specific immunological abnormalities that can explain the unique susceptibility to particular infections seen in AD-HIES have not yet been clarified.[34, 35]

Single-nucleotide polymorphism (SNPs) refers to single base differences in DNA among individuals. One of the interests in association studies is the association between SNPs and disease development.[36] The aim of this study, to identify functional SNPs within dbSNP located in coding regions of *STAT3* gene. This is the first study which covers an extensive in silico analysis of nsSNPs *STAT3* protein. This study will opens new doors to facilitate the development of novel diagnostic markers for associated diseases.[37–39]

## 2. Materials and Methods

### Data mining

The data on human *STAT3* gene was collected from National Center for Biological Information (NCBI) web site.[40] The SNP information (protein accession number and SNP ID) of the *PRSS1* gene was retrieved from the NCBI dbSNP (http://www.ncbi.nlm.nih.gov/snp/) and the protein sequence was collected from NCBI protein database (NCBI Reference Sequence: XP_024306664.1) (https://www.ncbi.nlm.nih.gov/protein/).

### SIFT

We used SIFT to observe the effect of A.A. substitution on protein function. SIFT predicts damaging SNPs on the basis of the degree of conserved amino A.A. residues in aligned sequences to the closely related sequences, gathered through PSI-BLAST.[41] It’s available at (http://sift.jcvi.org/).

### PolyPhen

PolyPhen (version 2) stands for polymorphism phenotyping version 2. We used PolyPhen to study probable impacts of A.A. substitution on structural and functional properties of the protein by considering physical and comparative approaches.[42] It’s available at (http://genetics.bwh.harvard.edu/pp2).

### Provean

Provean is an online tool that predicts whether an amino acid substitution has an impact on the biological function of a protein grounded on the alignment-based score. The score measures the change in sequence similarity of a query sequence to a protein sequence homolog between without and with an amino acid variation of the query sequence. If the PROVEAN score ≤−2.5, the protein variant is predicted to have a “deleterious” effect, while if the PROVEAN score is >−2.5, the variant is predicted to have a “neutral” effect.[43] It is available at (https://rostlab.org/services/snap2web/).

### Provean

Provean is an online tool that predicts whether an amino acid substitution has an impact on the biological function of a protein grounded on the alignment-based score. The score measures the change in sequence similarity of a query sequence to a protein sequence homolog between without and with an amino acid variation of the query sequence. If the PROVEAN score ≤−2.5, the protein variant is predicted to have a “deleterious” effect, while if the PROVEAN score is >− 2.5, the variant is predicted to have a “neutral” effect.[43] It is available at (https://rostlab.org/services/snap2web/).

### SNAP2

Functional effects of mutations are predicted with SNAP2 (29). SNAP2 is a trained classifier that is based on a machine learning device called "neural network". It distinguishes between effect and neutral variants/non-synonymous SNPs by taking a variety of sequence and variant features into account. The most important input signal for the prediction is the evolutionary information taken from an automatically generated multiple sequence alignment. Also structural features such as predicted secondary structure and solvent accessibility are considered. If available also annotation (i.e. known functional residues, pattern, regions) of the sequence or close homologs are pulled in. In a cross-validation over 100,000 experimentally annotated variants, SNAP2 reached sustained two-state accuracy (effect/neutral) of 82% (at anAUC of 0.9). In our hands this constitutes an important and significant improvement over othermethods.[44] It is available at (https://rostlab.org/services/snap2web/).

### SNPs&GO

Single Nucleotide Polymorphism Database (SNPs) & Gene Ontology (GO) is a support vector machine (SVM) based on the method to accurately predict the disease related mutations from protein sequence. FASTA sequence of whole protein is considered to be an input option and output will be the prediction results based on the discrimination among disease related and neutral variations of protein sequence. The probability score higher than 0.5 reveals the disease related effect of mutation on the parent protein function.[45] it’s available at (https://rostlab.org/services/snap2web/).

### PHD-SNP

An online Support Vector Machine (SVM) based classifier, is optimized to predict if a given single point protein mutation can be classified as disease-related or as a neutral polymorphism. It’s available at: (http://snps.biofold.org/phd-snp/phdsnp.html)

### I-Mutant 3.0

Change in protein stability disturbs both protein structure and protein function. I-Mutant is a suite of support vector machine, based predictors integrated in a unique web server. It offers the opportunity to predict the protein stability changes upon single-site mutations. From the protein structure or sequence. The FASTA sequence of protein retrieved from UniProt is used as an input to predict the mutational effect on protein and stability RI value (reliability index) computed.[46] It’s available at (http://gpcr2.biocomp.unibo.it/cgi/predictors/I-Mutant3.0/I-Mutant3.0.cgi).

### MUpro

MUpro is a support vector machine-based tool for the prediction of protein stability changes upon nonsynonymous SNPs. The value of the energy change is predicted, and a confidence score between −1 and 1 for measuring the confidence of the prediction is calculated. A score <0 means the variant decreases the protein stability; conversely, a score >0 means the variant increases the protein stability.[47] It’s available at (http://mupro.proteomics.ics.uci.edu/)

### I-Mutant 3.0

Change in protein stability disturbs both protein structure and protein function. I-Mutant is a suite of support vector machine, based predictors integrated in a unique web server. It offers the opportunity to predict the protein stability changes upon single-site mutations. From the protein structure or sequence. The FASTA sequence of protein retrieved from UniProt is used as an input to predict the mutational effect on protein and stability RI value (reliability index) computed.[46] It’s available at (http://gpcr2.biocomp.unibo.it/cgi/predictors/I-Mutant3.0/I-Mutant3.0.cgi).

### MUpro

MUpro is a support vector machine-based tool for the prediction of protein stability changes upon nonsynonymous SNPs. The value of the energy change is predicted, and a confidence score between −1 and 1 for measuring the confidence of the prediction is calculated. A score <0 means the variant decreases the protein stability; conversely, a score >0 means the variant increases the protein stability.[47] It’s available at (http://mupro.proteomics.ics.uci.edu/).

### GeneMANIA

We submitted genes and selected from a list of data sets that they wish to query. GeneMANIA approach to know protein function prediction integrate multiple genomics and proteomics data sources to make inferences about the function of unknown proteins.[48] It is available at (http://www.genemania.org/)

### Structural Analysis

#### Detection of nsSNPs Location in Protein Structure

Mutation3D is a functional prediction and visualization tool for studying the spatial arrangement of amino acid substitutions on protein models and structures. Mutation3D is able to separate functional from nonfunctional mutations by analyzing a combination of 8,869 known inherited disease mutations and 2,004 SNPs overlaid together upon the same sets of crystal structures and homology models. Further, it presents a systematic analysis of wholegenome and whole-exome cancer datasets to demonstrate that mutation3D identifies many known cancer genes as well as previously underexplored target genes.[49] It is available at (http://mutation3d.org).

#### Developing 3D structure of mutant STAT3 gene

The 3D structure of human Signal transducer and activator of transcription 3 (*STAT3*) protein is not available in the Protein Data Bank. Hence, we used RaptorX to generate a 3D structural model for wild-type *STAT3.* RaptorX is a web server predicting structure property of a protein sequence without using any templates..[50] It is available at (http://raptorx.uchicago.edu/).

#### Developing 3D structure of mutant STAT3 gene

The 3D structure of human Signal transducer and activator of transcription 3 (*STAT3*) protein is not available in the Protein Data Bank. Hence, we used RaptorX to generate a 3D structural model for wild-type *STAT3.* RaptorX is a web server predicting structure property of a protein sequence without using any templates..[50] It is available at (http://raptorx.uchicago.edu/).

#### Modeling Amino Acid Substitution

UCSF Chimera is a highly extensible program for interactive visualization and analysis of molecular structures and related data, including density maps, supramolecular assemblies, sequence alignments, docking results, conformational analysis[51] Chimera (version 1.8) available at (http://www.cgl.ucsf.edu/chimera/).

**Figure 1:**
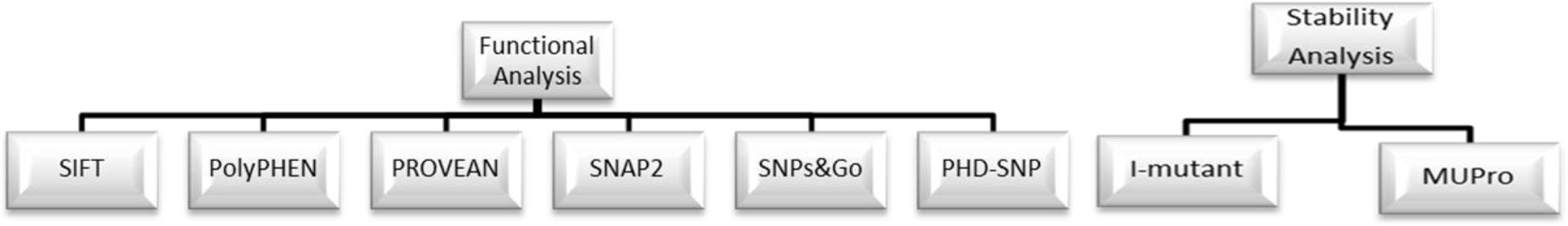
Diagrammatic representation for *STAT3* gene in coding region in silico work flow.

## Results

**Table1:** Damaging or Deleterious nsSNPs associated variations predicted by various softwares.

| dbSNP rs# | Sub | SIFT Prediction | Score | Polyphen Prediction | Score | PROVEAN Prediction | Score | SNAP2 Prediction | Score |
| --- | --- | --- | --- | --- | --- | --- | --- | --- | --- |
| rs869312892 | G716M | DAMAGING | 0 | possibly damaging | 0.518 | Deleterious | -7.28 | effect | 56 |
| rs1173682056 | P701R | DAMAGING | 0 | probably damaging | 0.98 | Deleterious | -8.119 | effect | 28 |
| rs1425974175 | I697G | DAMAGING | 0 | probably damaging | 1 | Deleterious | -5.12 | effect | 38 |
| rs1064796922 | E648K | DAMAGING | 0 | probably damaging | 1 | Deleterious | -2.741 | effect | 20 |
| rs1466712110 | F638K | DAMAGING | 0 | possibly damaging | 0.665 | Deleterious | -8.666 | effect | 19 |
| rs193922720 | W594K | DAMAGING | 0 | possibly damaging | 0.715 | Deleterious | -12.625 | effect | 74 |
| rs1064796762 | C582K | DAMAGING | 0.02 | possibly damaging | 0.714 | Deleterious | -7.764 | effect | 25 |
| rs781353111 | W542Y | DAMAGING | 0 | probably damaging | 0.996 | Deleterious | -8.32 | effect | 19 |
| rs776053336 | F524K | DAMAGING | 0 | probably damaging | 0.995 | Deleterious | -7.846 | effect | 29 |
| rs1281575473 | L510R | DAMAGING | 0 | possibly damaging | 0.818 | Deleterious | -4.957 | effect | 38 |
| rs145786768 | A507F | DAMAGING | 0 | possibly damaging | 0.649 | Deleterious | -5.355 | effect | 18 |
| rs763944667 | P503H | DAMAGING | 0.01 | possibly damaging | 0.855 | Deleterious | -5.908 | effect | 6 |
| rs146620441 | N498V | DAMAGING | 0 | probably damaging | 0.982 | Deleterious | -7.597 | effect | 34 |
| rs1442343135 | E407A | DAMAGING | 0.17 | probably damaging | 0.979 | Deleterious | -4.395 | effect | 14 |
| rs1401418993 | K392E | DAMAGING | 0.01 | probably damaging | 0.997 | Deleterious | -3.426 | effect | 29 |
| rs113994136 | R382Q | DAMAGING | 0 | possibly damaging | 0.939 | Deleterious | -3.355 | effect | 19 |
| - | R382L | DAMAGING | 0 | probably damaging | 0.975 | Deleterious | -5.941 | effect | 41 |
| rs113994135 | R382W | DAMAGING | 0 | probably damaging | 0.999 | Deleterious | -6.631 | effect | 68 |
| rs755524497 | R350M | DAMAGING | 0.23 | possibly damaging | 0.871 | Deleterious | -5.179 | effect | 47 |
| rs776115471 | R335Q | DAMAGING | 0 | probably damaging | 0.999 | Deleterious | -2.797 | effect | 3 |
| rs1427784630 | P330R | DAMAGING | 0 | probably damaging | 0.968 | Deleterious | -6.368 | effect | 12 |
| rs757347742 | Q248P | DAMAGING | 0.18 | probably damaging | 0.979 | Deleterious | -4.461 | effect | 2 |
| rs964892419 | R114C | DAMAGING | 0.05 | possibly damaging | 0.508 | Deleterious | -4.577 | effect | 14 |
| rs780604324 | C108Y | DAMAGING | 0.05 | possibly damaging | 0.92 | Deleterious | -3.748 | effect | 46 |

**Table2:** Disease effect nsSNPs associated variations predicted by SNPs&GO and PHD-SNP softwares.

| dbSNP rs# | Sub | SNPs&GO |  |  | PHD-SNP |  |  |
| --- | --- | --- | --- | --- | --- | --- | --- |
|  |  | Prediction | RI | Probability | Prediction | RI | Probability |
| rs1173682056 | P701R | Disease | 6 | 0.812 | Disease | 7 | 0.832 |
| rs1425974175 | I697G | Disease | 4 | 0.682 | Disease | 7 | 0.863 |
| rs193922720 | W594K | Disease | 8 | 0.923 | Disease | 9 | 0.964 |
| rs1064796762 | C582K | Disease | 6 | 0.801 | Disease | 7 | 0.846 |
| rs781353111 | W542Y | Disease | 4 | 0.687 | Disease | 4 | 0.686 |
| rs776053336 | F524K | Disease | 7 | 0.841 | Disease | 8 | 0.875 |
| rs1281575473 | L510R | Disease | 4 | 0.678 | Disease | 7 | 0.85 |
| rs145786768 | A507F | Disease | 6 | 0.797 | Disease | 8 | 0.901 |
| rs763944667 | P503H | Disease | 3 | 0.665 | Disease | 4 | 0.718 |
| rs146620441 | N498V | Disease | 6 | 0.777 | Disease | 5 | 0.768 |
| rs780604324 | C108Y | Disease | 0 | 0.522 | Disease | 6 | 0.824 |

**\*RI: Reliability Index**

**Table3:**
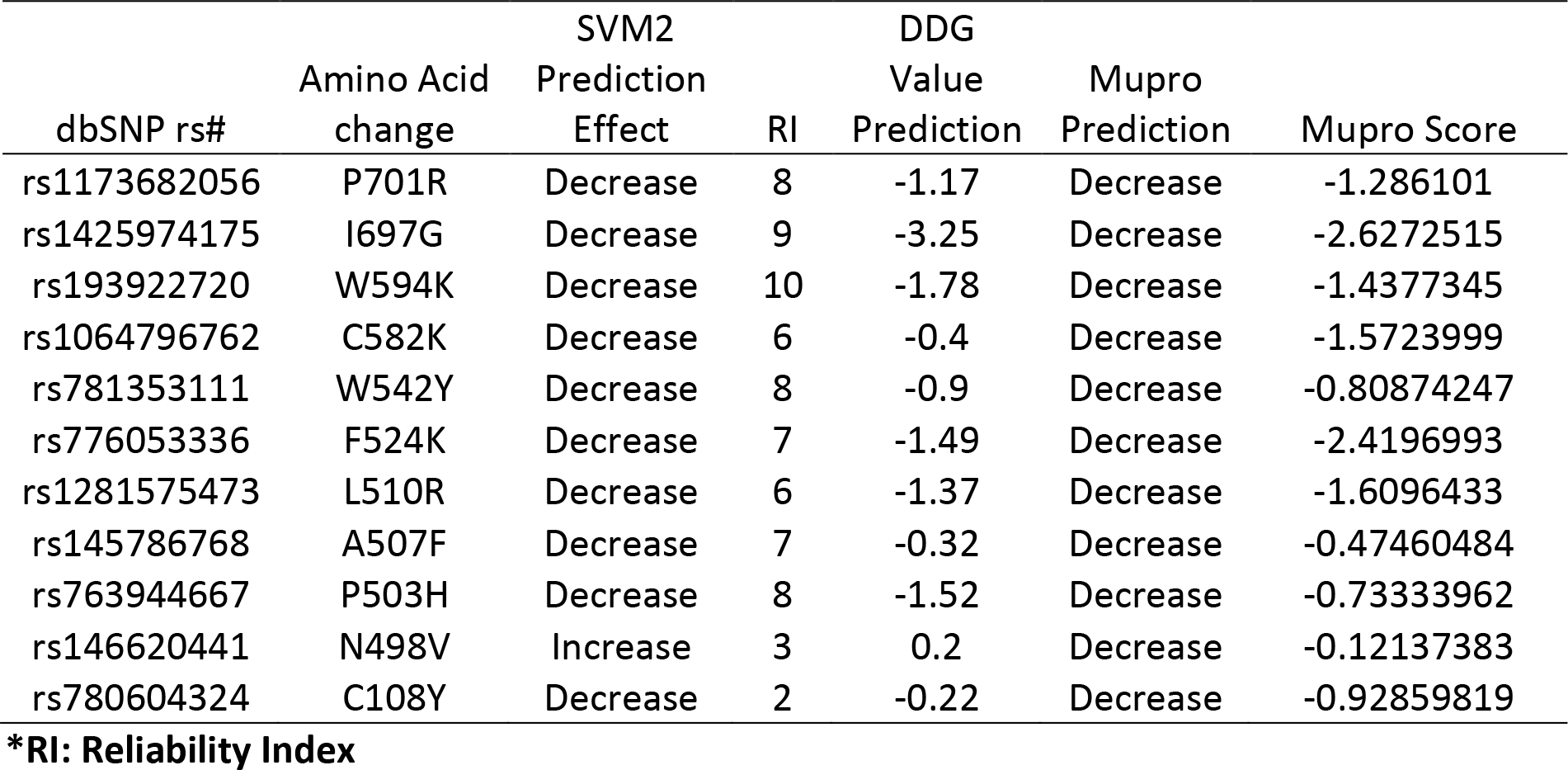
Stability analysis predicted by I-Mutant version 3.0 and MUPro (also Show the 11 novel mutations)

**Table 4:** The *STAT3* gene functions and its appearance in network and genome

| Function | FDR | Genes<br>in<br>network | Genes<br>in<br>genome |
| --- | --- | --- | --- |
| cellular response to growth hormone stimulus | 0.011108 | 3 | 31 |
| response to peptide | 0.011108 | 5 | 260 |
| cellular response to peptide | 0.011108 | 5 | 247 |
| tumor necrosis factor-mediated signaling pathway | 0.011108 | 3 | 35 |
| growth hormone receptor signaling pathway | 0.011108 | 3 | 31 |
| response to growth hormone | 0.011108 | 3 | 32 |
| response to estradiol | 0.011108 | 3 | 24 |
| JAK-STAT cascade | 0.011108 | 4 | 91 |
| response to peptide hormone | 0.011108 | 5 | 255 |
| JAK-STAT cascade involved in growth hormone signaling pathway | 0.011108 | 3 | 24 |
| response to organic cyclic compound | 0.011108 | 5 | 246 |
| cellular response to peptide hormone stimulus | 0.011108 | 5 | 244 |
| regulation of peptidyl-tyrosine phosphorylation | 0.019403 | 4 | 138 |
| response to estrogen | 0.021834 | 3 | 47 |
| positive regulation of response to DNA damage stimulus | 0.021834 | 3 | 47 |
| regulation of protein complex assembly | 0.023675 | 4 | 153 |
| response to alcohol | 0.063704 | 3 | 75 |
| peptidyl-tyrosine phosphorylation | 0.063704 | 4 | 210 |
| cellular response to epidermal growth factor stimulus | 0.063704 | 2 | 11 |
| peptidyl-tyrosine modification | 0.063704 | 4 | 210 |
| cellular response to tumor necrosis factor | 0.063704 | 3 | 74 |
| positive regulation of ERK1 and ERK2 cascade | 0.077484 | 3 | 83 |
| temperature homeostasis | 0.077484 | 2 | 14 |
| astrocyte differentiation | 0.077484 | 2 | 14 |
| response to steroid hormone | 0.077484 | 3 | 82 |
| regulation of homotypic cell-cell adhesion | 0.079432 | 2 | 15 |
| positive regulation of protein complex assembly | 0.079432 | 3 | 90 |
| response to tumor necrosis factor | 0.079432 | 3 | 87 |
| response to epidermal growth factor | 0.079432 | 2 | 15 |
| response to lipid | 0.084124 | 4 | 249 |

**Table (5).**
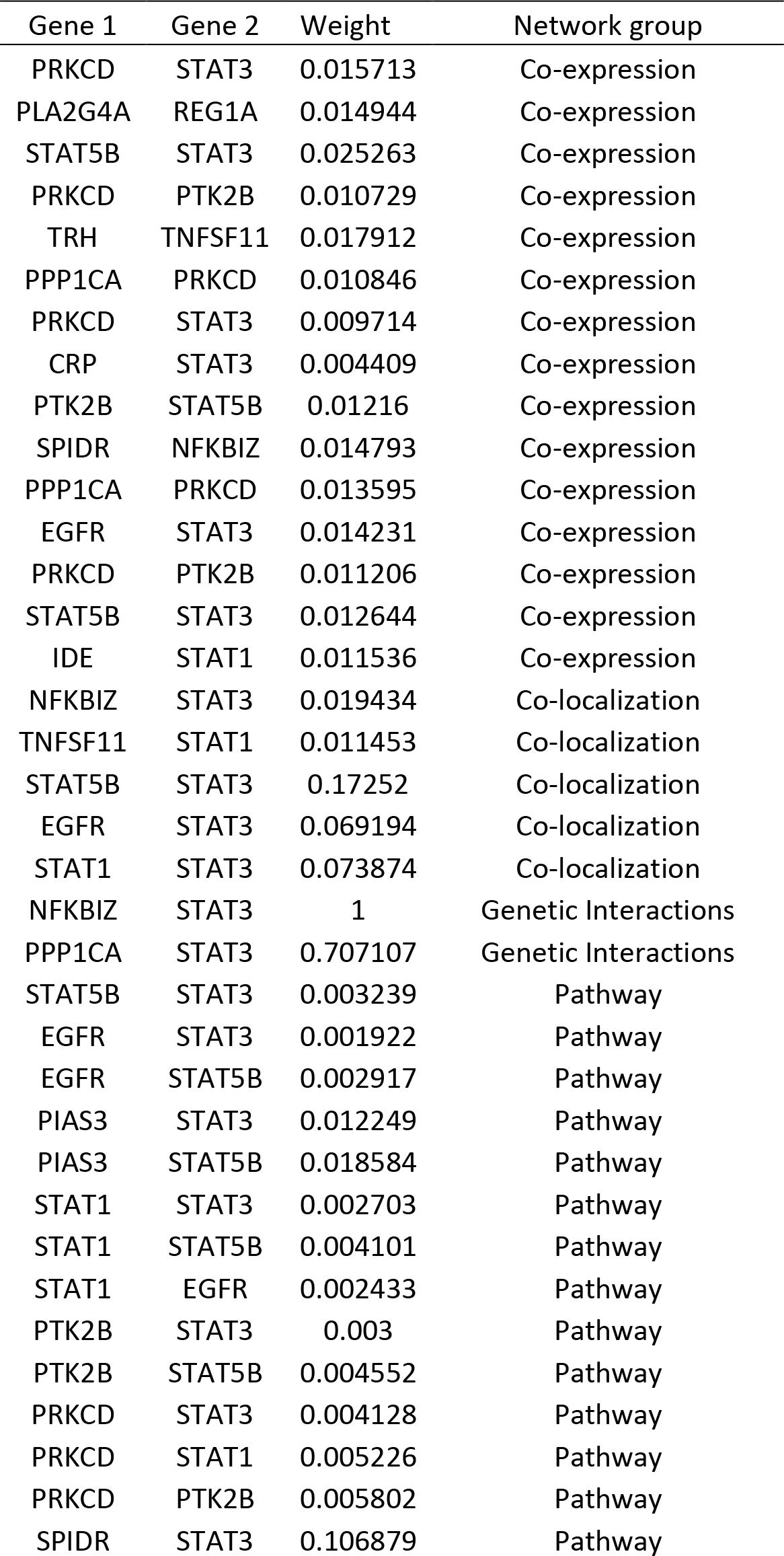
The gene co-expressed, share domain and Interaction with *STAT3* gene network

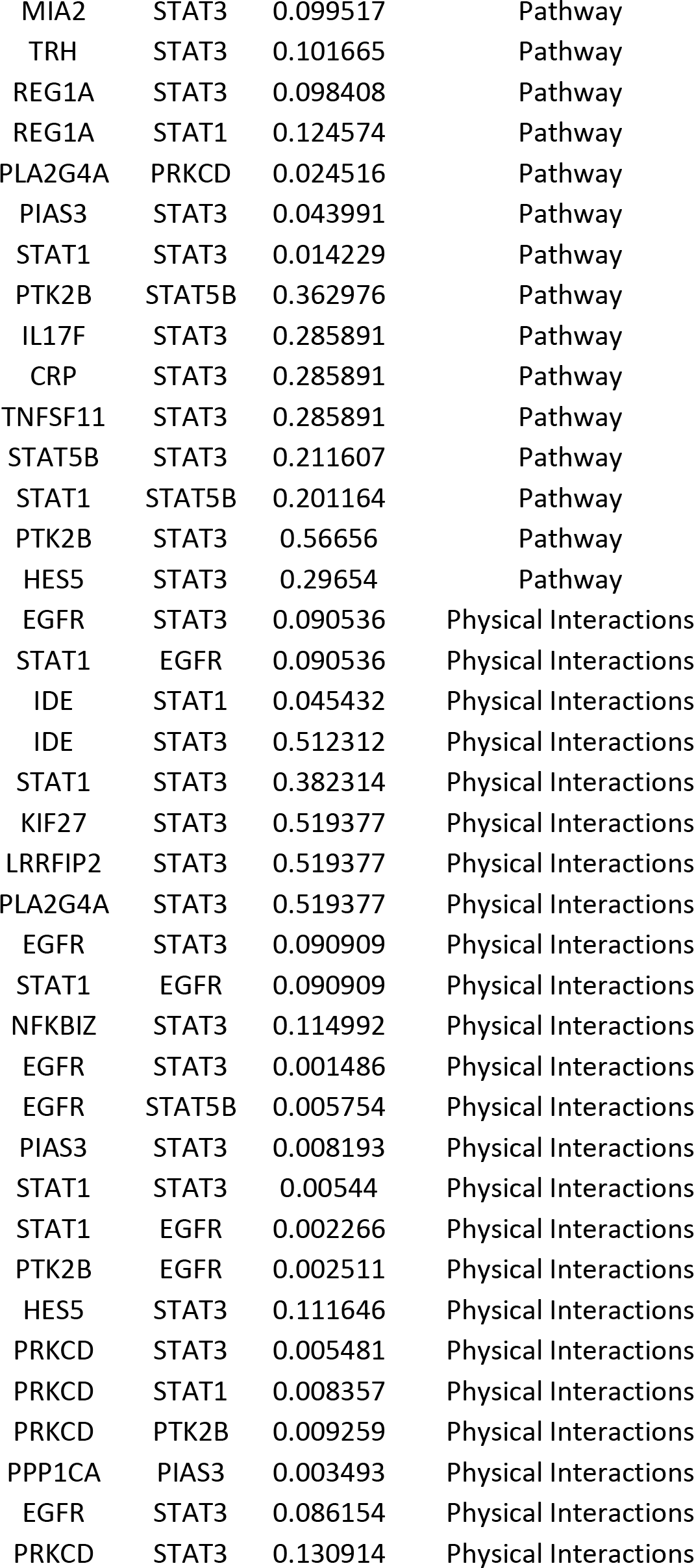

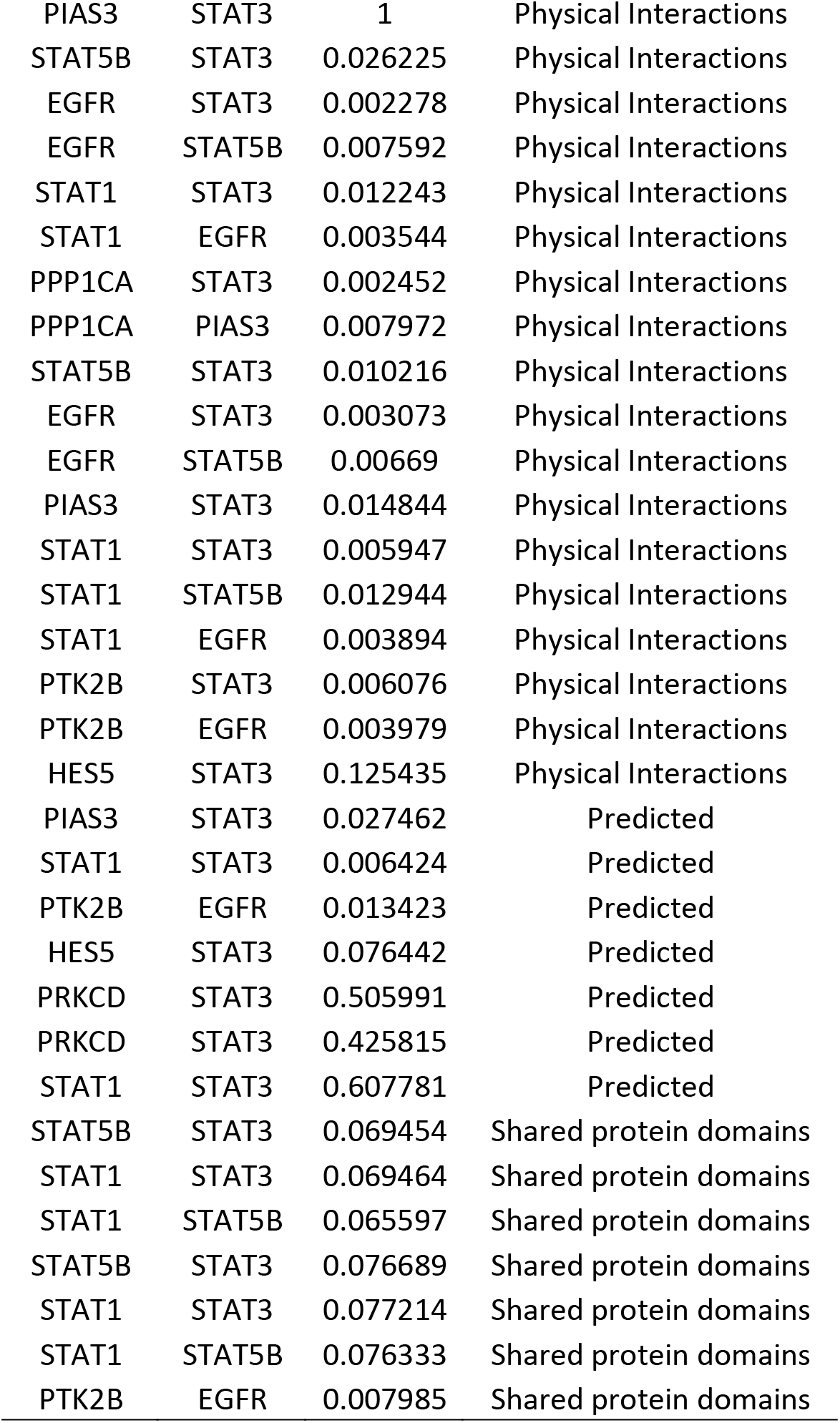

**Figure 2:**
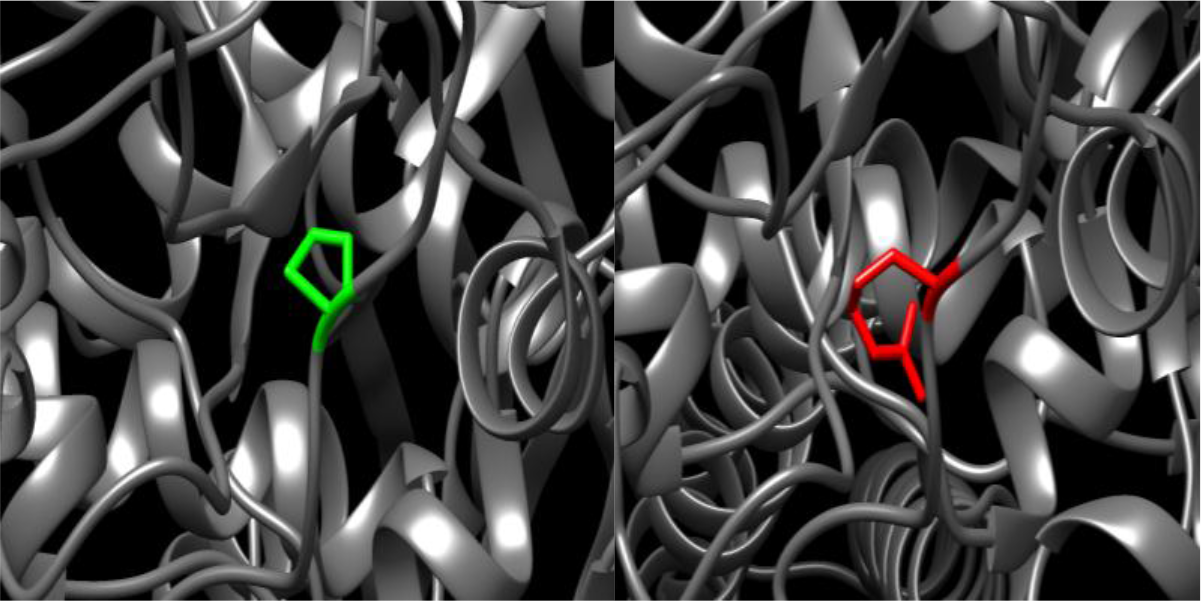
(P701R): change in the amino acid Proline into Arginine at position 701.

**Figure 3:**
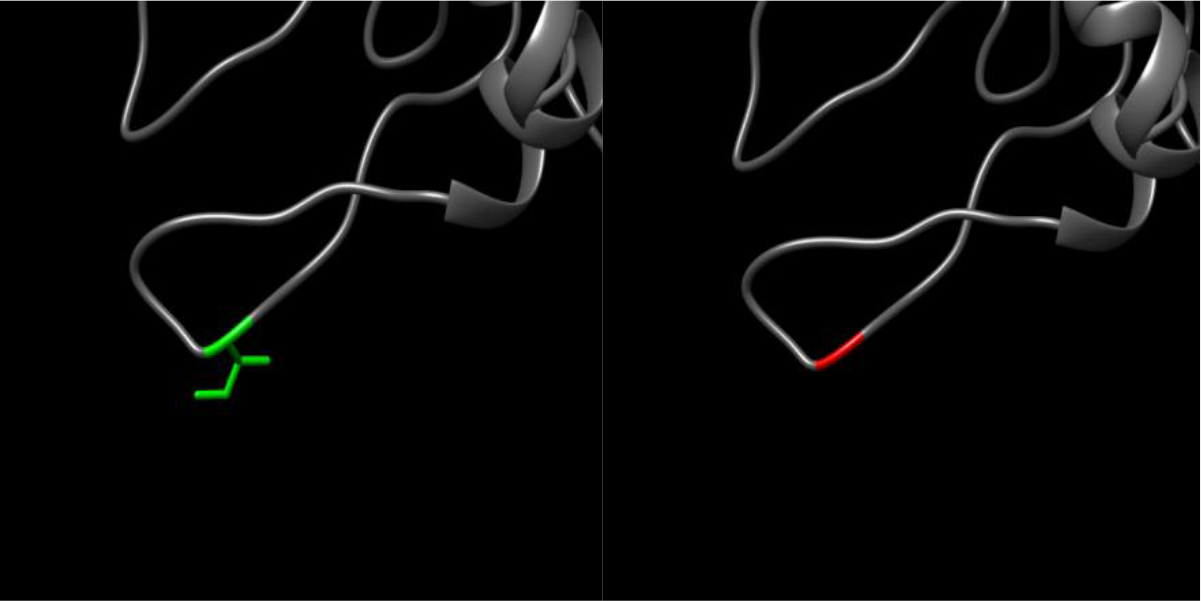
(I697G): change in the amino acid Proline into Arginine at position 701.

**Figure 4:**
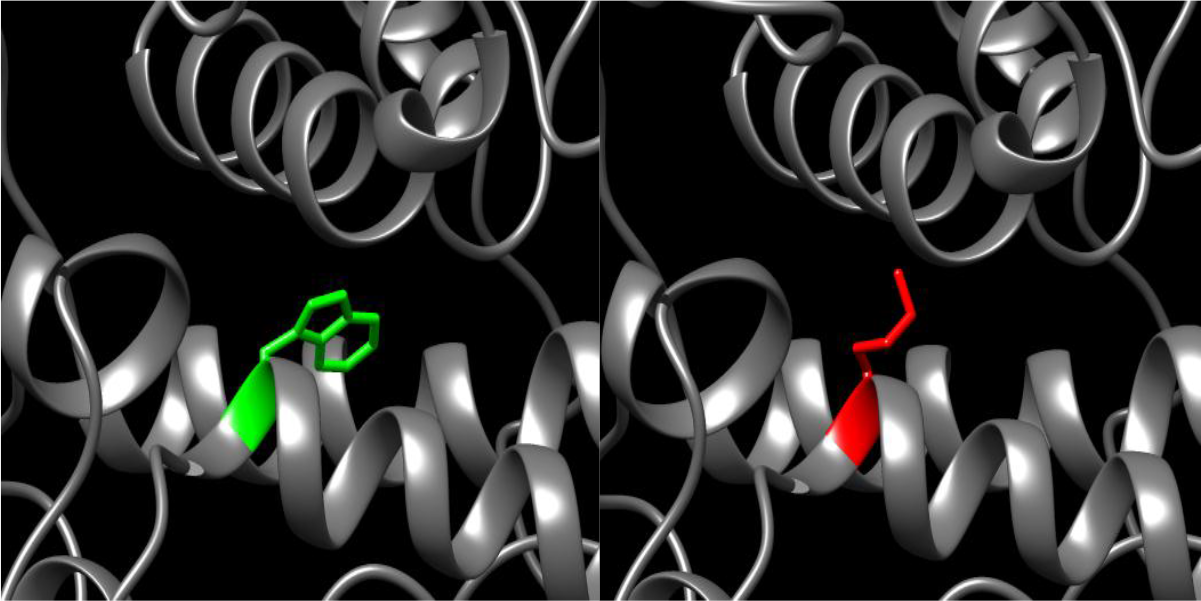
(W594K): change in the amino acid Tryptophan into Lysine at position 594.

**Figure 5:**
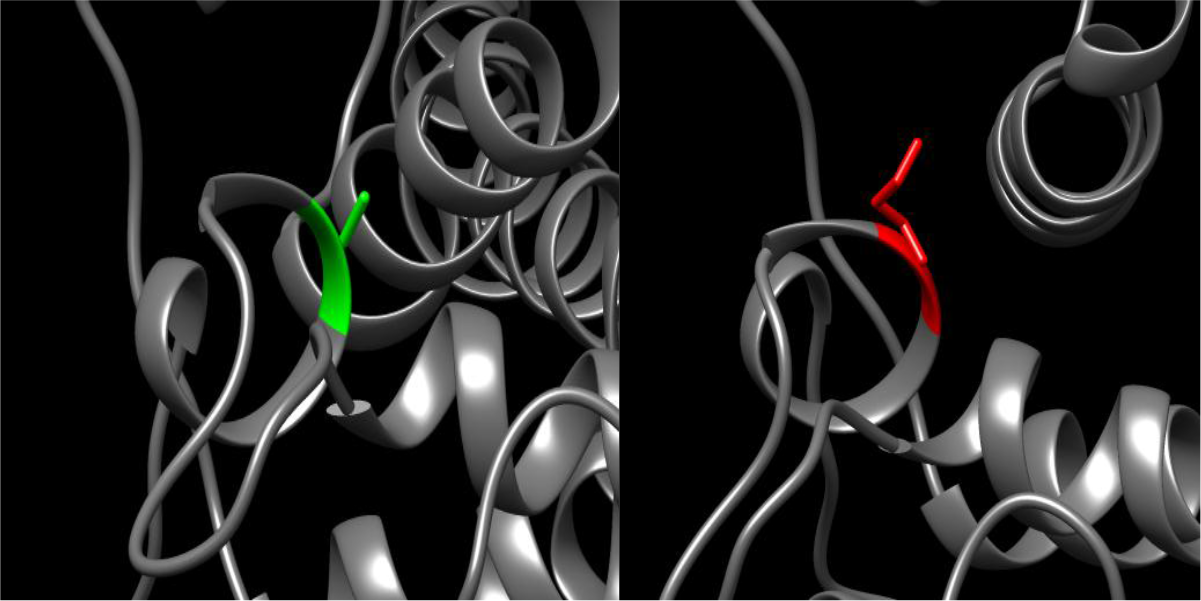
(C582K): change in the amino acid Cysteine into Lysine at position 594.

**Figure 6:**
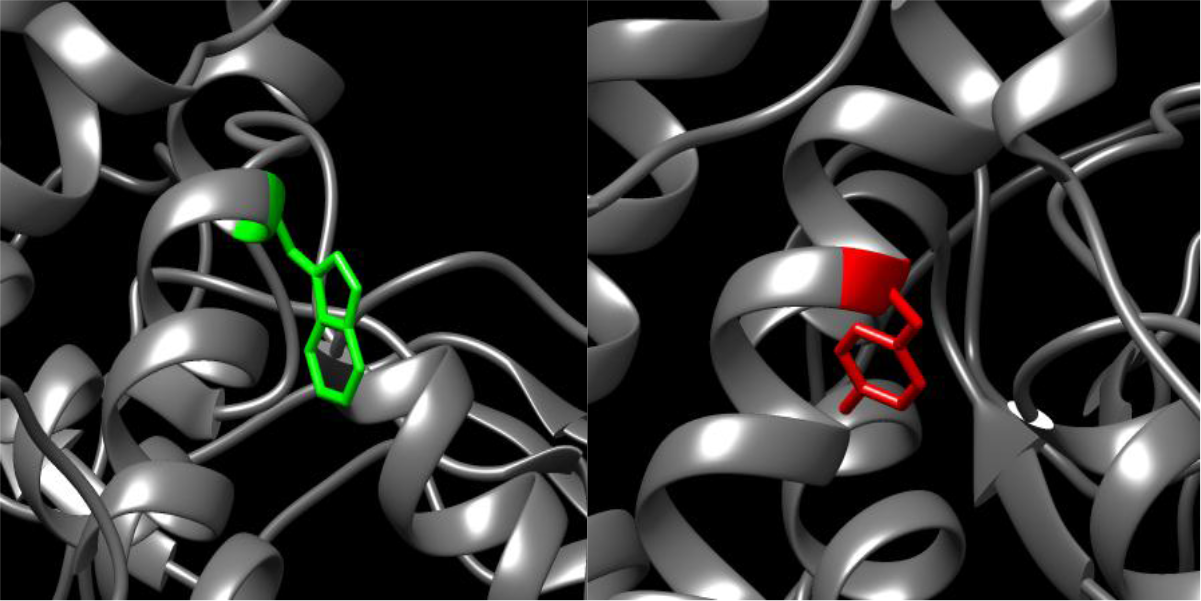
(W542Y): change in the amino acid Tryptophan into Tyrosine at position 542.

**Figure 7:**
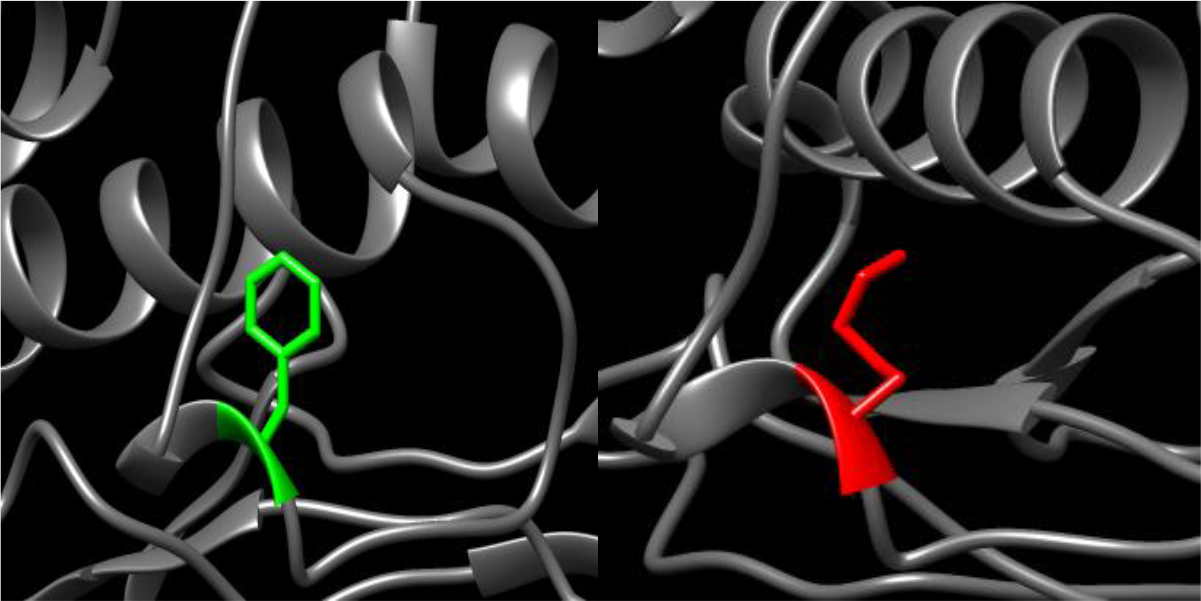
(F524K): change in the amino acid Phenylalanine into Lysine at position 542.

**Figure 8:**
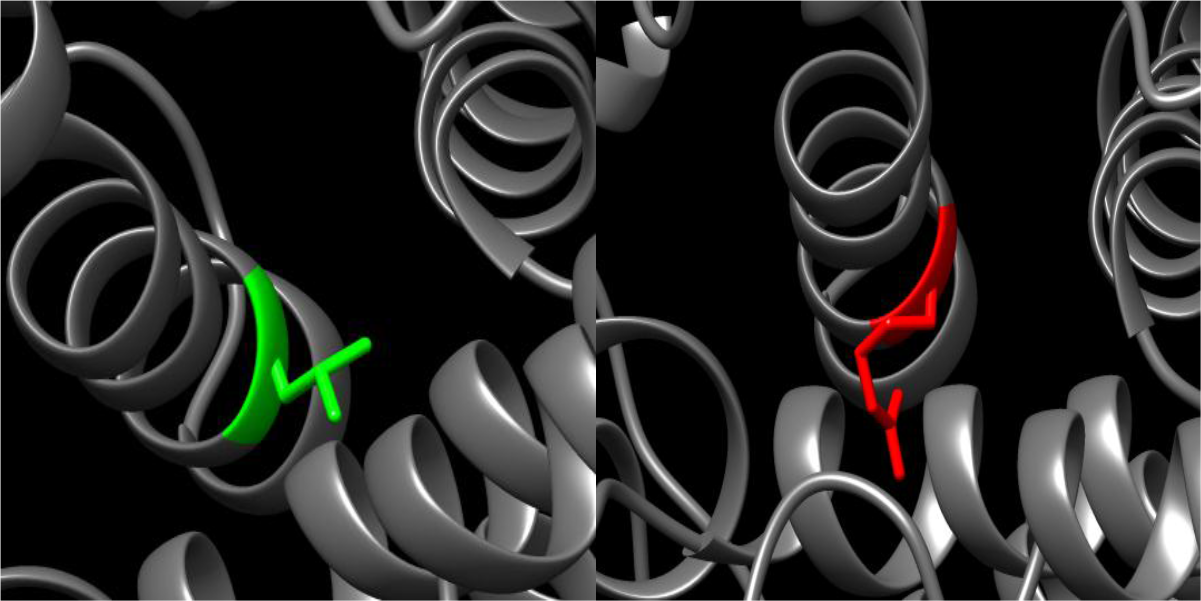
(L510R): change in the amino acid Leucine into Arginine at position 542.

**Figure 9:**
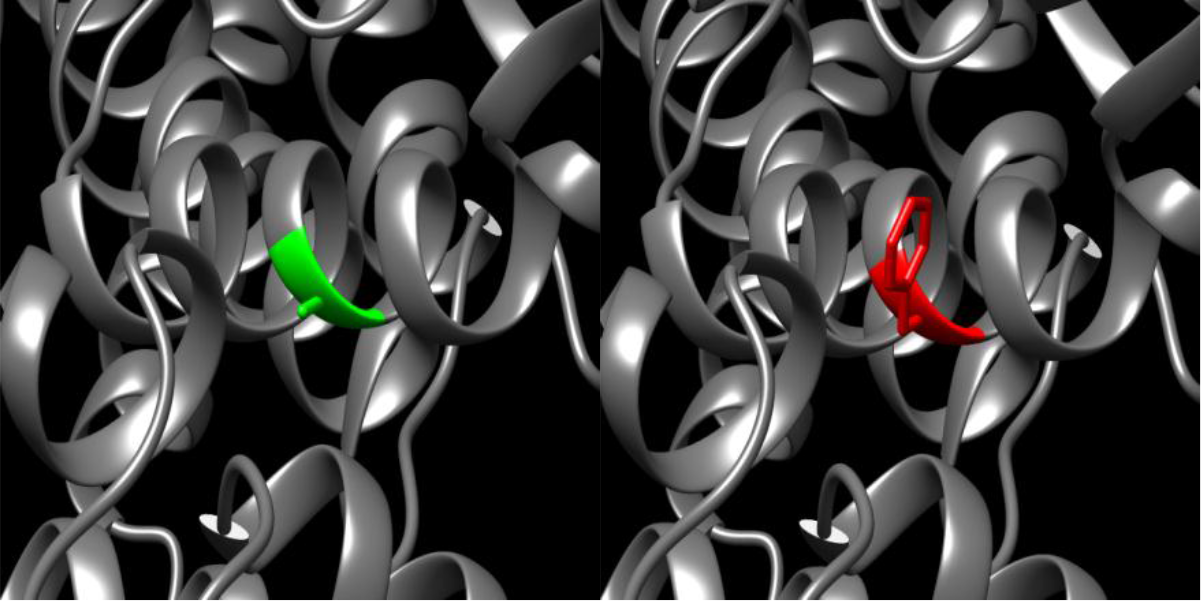
(A507F): change in the amino acid Alanine into Phenylalanine at position 507.

**Figure 10:**
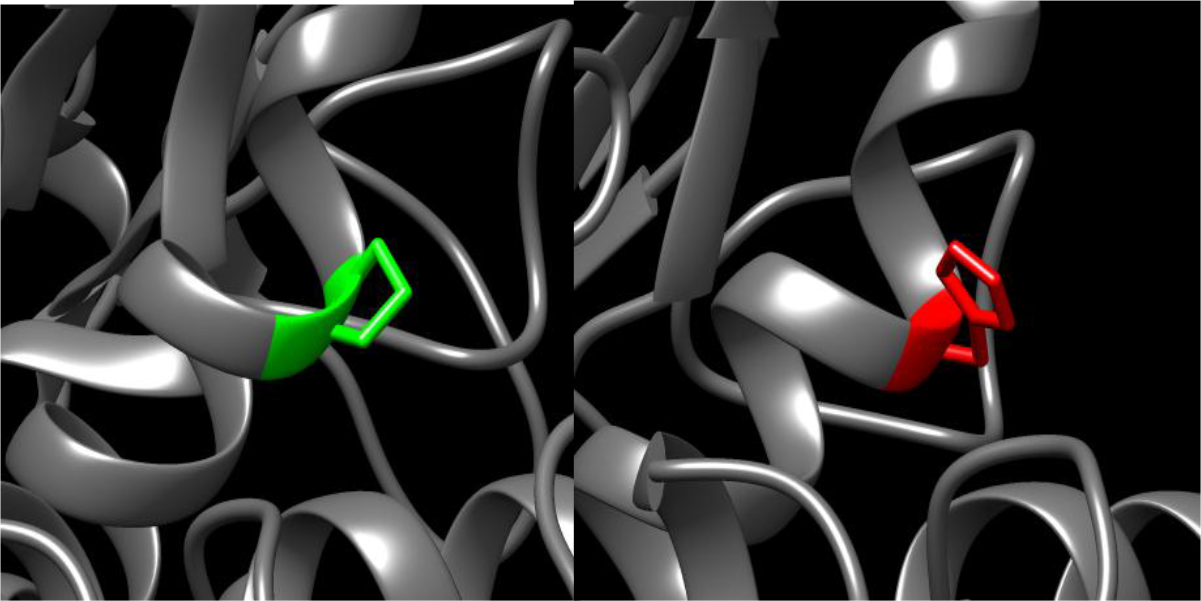
(P503H): change in the amino acid Proline into Histidine at position 542.

**Figure 11:**
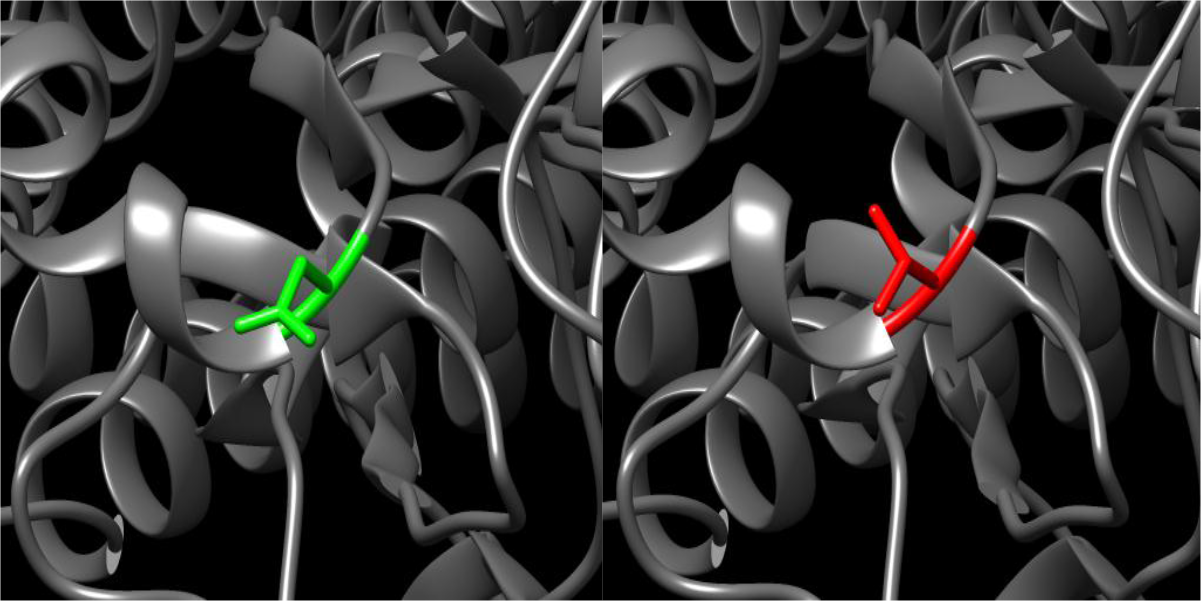
(N498V): change in the amino acid Asparagine into Valine at position 498.

**Figure 12:**
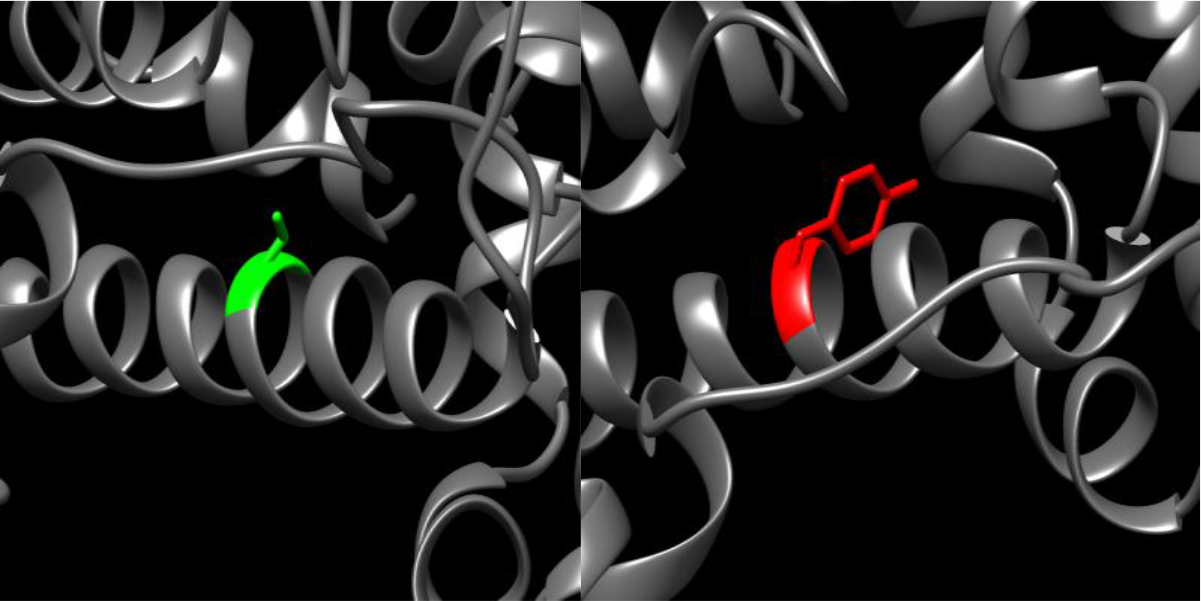
(C108Y): change in the amino acid Cysteine into Tyrosine at position 542.

**Figure 13:**
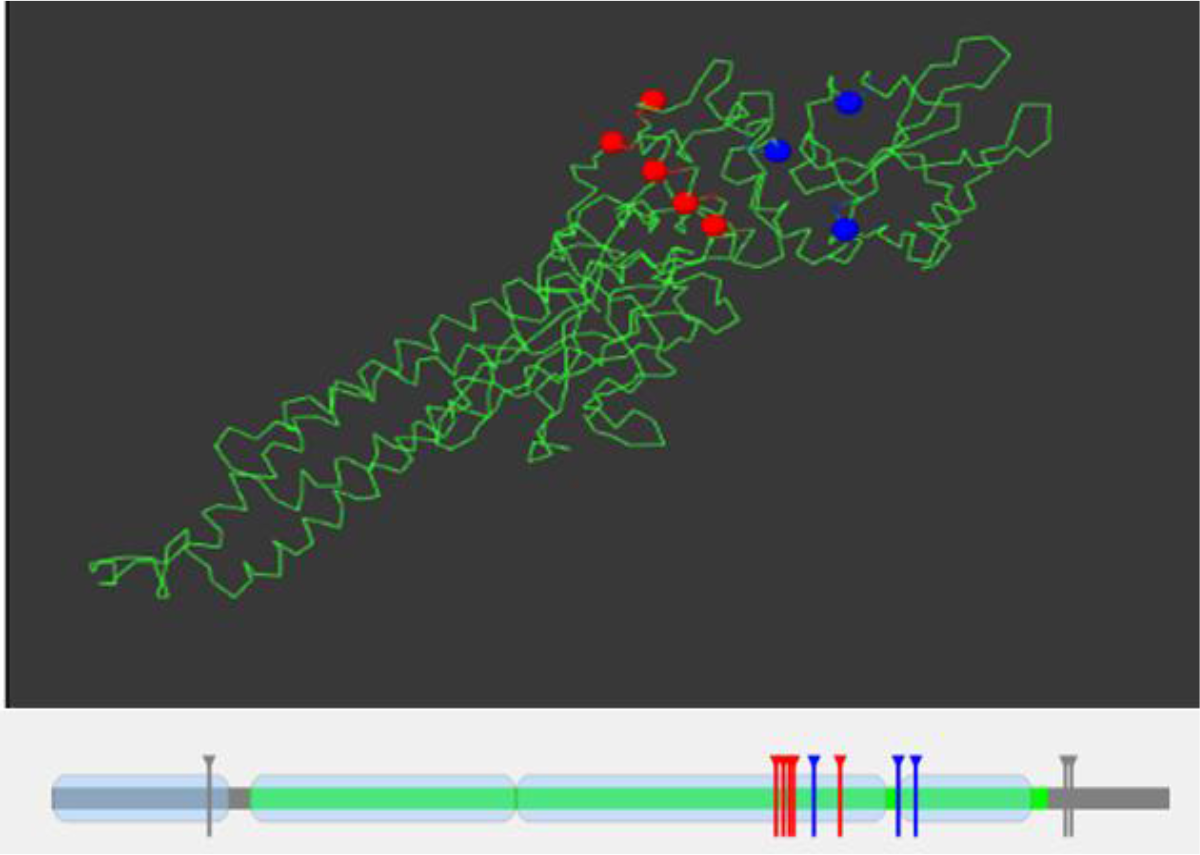
Structural models for wild type *STAT3*, illustrated by Mutation3D.

**Figure 14:**
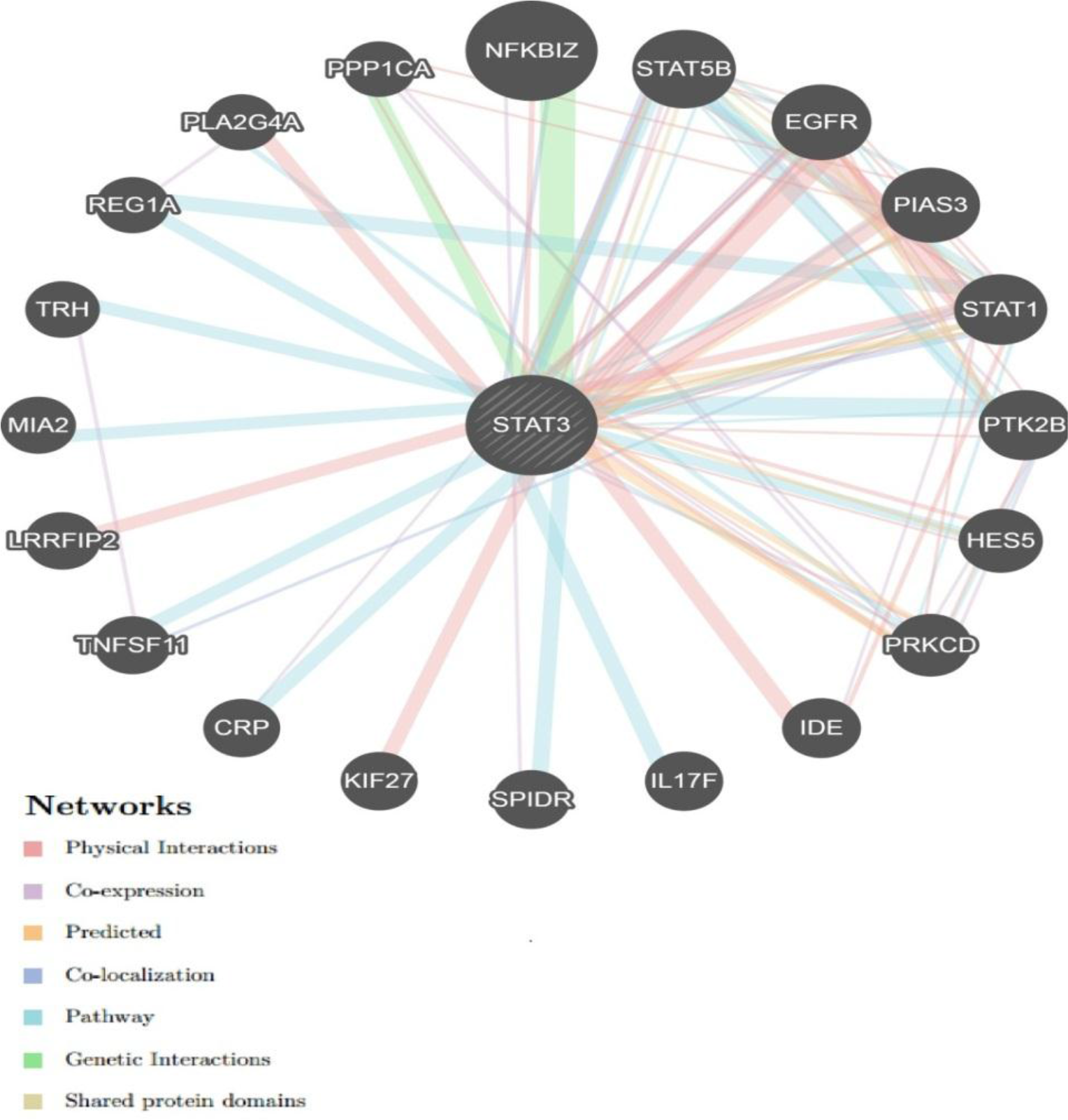
Interaction between *STAT3* and its related genes.

## Discussion

Eleven novel SNPs were found to be deleterious in this study. The in silico methods used to analyze the SNPs were based on different aspects and parameters describing the pathogenicity and provide clues on the molecular level about the effect of mutation on the structure and function of the final protein.(figure 1) 958 SNPs were retrieved form NCBI database, of which 251 Were found to be missense, we first analyze the deleterious effect of different candidate SNPs on the function of the final protein through 4 softwares (SIFT, POLYHEN2, PROVEAN and SNPA2)and we found that, the shared deleterious SNPs between them to be 24 SNPs.(table 1) we further analyses it through (SNPS&GO and PHD-SNP) and we found the double positive result to be 11 SNPs.(table 2)

Stability analysis which predicted by I-mutant 3.0 and MUPro servers revealed that, all SNPs had decreased protein stability by both servers except N498V (rs146620441) had predicted by I-mutant 3.0, increased protein stability. (Table 3) for further study the structural changes we used chimera software (figures 2 to 12), In the data extracted from the NCBI database the substitutions (R382W) (rs113994135), V637M (rs113994139) were found to be pathogenic which correspond to what is mentioned in other paper. [53] The substitution R382W (rs113994135) was also found to be deleterious by four soft wares in this study which correspond to previous studies.[53, 54]

GeneMANIA revealed that *STAT3* has many vital functions: blood microparticle, cobalamin metabolic process, extracellular matrix organization, extracellular structure organization, serine hydrolase activity, serine-type endopeptidase activity, serine-type peptidase activity. The genes co-expressed with, share similar protein domain, or participate to achieve similar function were illustrated by GeneMANIA and shown in figure (14) Tables (4 & 5). Additionally, we performed analysis by Mutation3D, our result show that: (N498V, P503H, A507F, L510R and F524K) located in the domain. (Figure 13)

The STAT3 gene is associated with other disease like inflammatory bowel disease along other genes like IL23R and JAK2 genes.[55] there are also a relations with Autoimmune lymphoproliferative syndrome (ALPS) mainly in p.R278H, p.M394T SNPs,[56] STAT3 also becomes persistently activated in a high percentage of malignancies (e.g. breast, prostate, ovarian, and colon cancers), thus contributing to malignant transformation and progression which makes STAT3 an attractive therapeutic target for cancers.[57]there is also an evidence that studying gene expression of STAT1, STAT2 and STAT3 gene can be useful for evaluating the efficacy of IFN treatment of the MS patients. [58]

Our study is the first in silico analysis which based on functional and structural analysis while all other previous studies based on in vivo analysis, molecular analysis and genome sequencing [53, 59–61] This study revealed eleven Novel Pathological mutations have a potential functional impact and may thus be used as diagnostic markers for Job’s syndrome. Finally some appreciations of wet lab techniques are suggested to support our findings.

## Conclusion

A total of eleven novel nsSNPs were predicted to be responsible for the structural and functional modifications of *STAT3* protein. The newly recognized genetic cause of the Autosomal dominant hyper-IgE syndrome affects complex, compartmentalized somatic and immune regulation. This study will opens new doors to facilitate the development of novel diagnostic markers for associated diseases.

## Acknowledgment

The authors wish to acknowledgment the enthusiastic cooperation of Africa City of Technology - Sudan.

